# Synthetic data generation with probabilistic Bayesian Networks

**DOI:** 10.1101/2020.06.14.151084

**Authors:** Grigoriy Gogoshin, Sergio Branciamore, Andrei S. Rodin

## Abstract

Bayesian Network (BN) modeling is a prominent and increasingly popular computational systems biology method. It aims to construct probabilistic networks from the large heterogeneous biological datasets that reflect the underlying networks of biological relationships. Currently, a variety of strategies exist for evaluating BN methodology performance, ranging from utilizing artificial benchmark datasets and models, to specialized biological benchmark datasets, to simulation studies that generate synthetic data from predefined network models. The latter is arguably the most comprehensive approach; however, existing implementations are typically limited by their reliance on the SEM (structural equation modeling) framework, which includes many explicit and implicit assumptions that may be unrealistic in a typical biological data analysis scenario. In this study, we develop an alternative, purely probabilistic, simulation framework that more appropriately fits with real biological data and biological network models. In conjunction, we also expand on our current understanding of the theoretical notions of causality and dependence / conditional independence in BNs and the Markov Blankets within.

## 1 Introduction

Dependency modeling is a descriptive / predictive modeling activity that is especially suited to a systems biology approach to data analysis (in particular, analysis of heterogeneous big data). During dependency modeling, a biological network, or a graphical representation of a biological model (genotypes, phenotypes, metabolites, endpoints, and other omics variables and their interrelationships), is constructed from the “flat” data. (Subsequently, specific components of biological networks can be evaluated using traditional / parametric statistical criteria, and sub-networks can be selected for further biological hypothesis generation and testing). This activity is also known as causal discovery (inference) in graphical models. Bayesian Networks (BNs)-based dependency modeling is an established computational biology tool that is rapidly gaining acceptance in biological big data analysis (Branciamore et al. (2018); Cooper et al. (2015); Gogoshin et al. (2017); Jiang et al. (2010); Lan et al. (2016); Neapolitan et al. (2014); Needham et al. (2007); Pe’er (2005); Piatetsky-Shapiro and Tamayo (2003); Qi et al. (2014); Rodin et al. (2005, 2012); Sedgewick et al. (2019); Sherif et al. (2015); Wang et al. (2019); Yin et al. (2015); Zeng et al. (2016); Ziebarth et al. (2013); Zhang and Shi (2017); Zhang et al. (2017, 2014)). Comprehensive treatments of BN methodology, and probabilistic networks (PNs) in general, are found in numerous textbooks (Pearl (1988, 2009); Russell and Norvig. (2009); Spirtes et al. (2000)) and reviews (Daly et al. (2011); Glymour et al. (2019); Heckerman et al. (1995); Heckerman (1995); Spirtes and Zhang (2016); Zhang et al. (2018)).

Most of the recent work in the field aimed at improving the scalability of BN modeling, handling mixed data types (including various types of biological data), incorporating latent variables, and developing more robust software interfaces and visualization options (Andrews et al. (2018, 2019); Chen et al. (2019); Hong et al. (2018); Jabbari et al. (2017); Ogarrio et al. (2016); Raghu et al. (2018); Ramsey et al. (2017); Sedgewick et al. (2016); Spirtes and Zhang (2016); Xing et al. (2017, 2018); Yu et al. (2019); Zhang et al. (2018, 2019)). In view of these developments, the ability to objectively assess the performance of BN modeling is of the utmost importance. Currently, there are three principal venues for accomplishing this: (i) using well-established (in machine learning community) predefined benchmark models/datasets, such as “Alarm” or “Asia”, (ii) using various specialized biologically-oriented benchmark datasets, both real and simulated, such as “DREAM” (Eicher et al. (2019); Wang et al. (2019); Xing et al. (2017)), and (iii) developing approximately realistic simulation frameworks (Andrews et al. (2018); Han et al. (2017); Ellis and Wong (2008); Isci et al. (2011); Tasaki et al. (2015); Zhang et al. (2019)). The first approach is necessarily limited and does not generalize to modern high-dimensional biological data. The latter two strategies have pros and cons; a useful discussion can be found in (Wang et al. (2019)). In this study, we concentrate on the third approach, namely constructing robust, generalizable, and mathematically rigorous simulation frameworks. Currently, these frameworks typically involve (i) specifying synthetic network structures, and (ii) utilizing linear SEM (structural equation modeling) methodology to generate synthetic data (Isci et al. (2011); Tasaki et al. (2015); Zhang et al. (2019)).

In the broader context of heterogeneous biological data and networks, it is important to distinguish between the notions of causality, correlation, association and dependence / conditional independence. Establishing causality (Cooper et al. (2015); Glymour et al. (2019); Pearl (2009); Zhang et al. (2018)) is usually perceived as the ultimate goal in the biological network analyses (Ramsey et al. (2017); Zeng et al. (2016); Zhang et al. (2019)). However, establishing (and even defining) causality is often equivocal, for both theoretical (Glymour et al. (2019); Zhang et al. (2018)) and practical/biological reasons (e.g., ambiguity of the “causal” interpretation of the relationship between two correlated gene expression measurements, or two single nucleotide polymorphisms in linkage disequilibrium) (Tasaki et al. (2015)). Of course, there are advantages to viewing certain biological network models (or their parts) through the lens of directional causality. First, directional causality often fits well to biological reality (to use an obvious example, phenotypes tend to be “downstream” of genotypes). Second, existing BN algorithmic machinery largely relies on the concept of directed acyclic graphs (DAGs). And yet, at the most basic level (e.g., physical / chemical interaction of two biological molecules), it is unclear if it is possible, or even desirable, to impose a “directional causation” label on what is essentially a dependence relationship that is inferred from observing the joint probability distribution. The concept of “cause - effect” may have more to do with rationalization in seeking implicational chains rather than reflecting the totality of any given interaction. Therefore, we chose to frame this study predominantly around the notion of dependence / conditional independence, which is a measure of relevance that allows us to make inferences about biological reality without invoking considerations of causality (however, we do account for ancestor/descendant relationships in DAGs when moving from general PNs to the BN implementation).

Regardless of whether we focus on causation or dependence relationships, SEM might not be the most appropriate tool for generating synthetic data in the case of biological BNs. Leaving aside the issues of acyclicity (and directionality in general), and the assumptions of linearity and normality (which can be dealt with, if imperfectly), there is a fundamental issue of observability. Indeed, consider a typical biological dataset subject to BN modeling — “real” biological datasets include only comparatively small sets of variables that are observable under the confines of specific biomedical studies (i.e., experiments). These variables cannot be expected to be in any particular categorizable relationship known or suspected a priori. The most one can postulate about a “real” dataset is that the variables in it may be dependent or independent, and certainly not that these variables belong to a carefully selected group of key components that adequately describe the dynamics of a biological system/model in question. Therefore, synthetic datasets generated via SEM are fundamentally different from real biological datasets. Instead, we propose to use PNs as the general abstraction for synthetic data generation purposes. Such an abstraction, built around an analytically defined distribution induced by a given graphical model and coupled with random model structure generation, is immediately interpretable as the joint probability distribution, while simultaneously featuring inferential characteristics encoded in conditional independencies. However, building a simulation framework around PNs in a mathematically rigorous fashion is not a trivial undertaking. Below we detail considerations involved in building such a framework in Methods, Proofs and Results: Sections 2–3; ascertain the congruence between the ‘‘forward” process (simulations) and ‘inverse” process (actual reconstruction from the data of the networks and Markov Blankets within) in Methods, Proofs and Results: Section 4; and discuss simulation sample size considerations in Results: Section 5.

## 2 Methods, Proofs and Results: Synthetic data generation with probabilistic networks

Synthetic data generation is a “forward problem” in which data is sampled from a distribution describing the model under consideration. Similarly, the “inverse problem” is BN modeling per se, i.e., reconstructing a BN model from the observed data.

Because we are predominantly interested in performance testing of probabilistic inference over Markov Blankets of particular variables in the network, as well as in assessing the quality of extraction of structural parameters of a model from the data induced by it, the data synthesis algorithm should be able to generate samples from arbitrarily complex models.

The analytically defined distribution induced by a given model has a factorizable joint probability that is no more complex than the basic chain rule. Given the set of further simplifications dictated by conditional independencies encoded in the model, the complexity of the factorization may be significantly lower than the worst case scenario.

In the most reduced form the chain factorization of the joint probability of a given model with an appropriate indexing of variables is

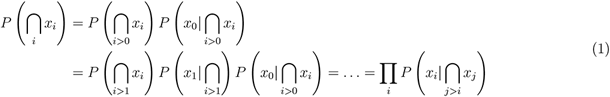

This product of conditional probabilities which represents the joint probability for the model provides a way to generate sample components serially from local conditional probabilities instead of direct sampling from the n-dimensional joint distribution. This is an important consideration, because using direct sampling to populate low probability fine event structures is difficult given a finite sample size. In fact, this difficulty grows with the dimensionality of the model, so that as the number of variables grows, many joint events quickly become intractable even for binary variables. Therefore, being able to rely on local sampling of conditional probability distributions is a key factor in successfully synthesizing components of an adequate n-dimensional sample that actually reflects the fine structure of the joint distribution. However, if care is not taken to obtain such a level of resolution in the joint distribution, then the resulting synthetic sampling is likely to run into source uniqueness problems. That is to say, the same data could be obtained from different source distributions, making the source distinction all but impossible.

In order to utilize the resolution provided by the expansion above, we need to find a way to methodically obtain conditional distributions for the dependent variables. Although we will only look at binary distributions, the results readily generalize to higher arity.

Consider elementary mechanics for discretely distributed variables — in the linear algebra sense, the joint distribution of two variables *x* and *y* can be represented as a matrix of joint events, and is readily decomposed by the chain law into an inner product of conditional distribution and unconditional distribution, as follows:

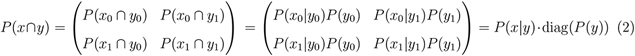

where individual events are notated as indexed variables and *P*(*y*) = (*P*(*y*_0_), *P*(*y*_1_))^*T*^.

The marginalization operation for the lexicographic order provided above is given by

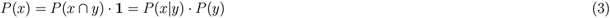

and

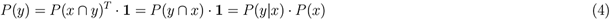

where **1** = (1,1)^*T*^. Note that here the change of the order of arguments in the joint probability operator implies the transposition operation, i.e. *P*(*x* ∩ *y*)^*T*^ = *P*(*y* ∩ *x*).

We can now derive the condition necessary for generating conditional distributions for *dependent* variables. A formal criterion for dependence is obtained by negation of the independence criterion

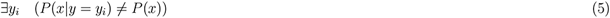

i.e. the existence of at least one dependent event is sufficient to establish dependence.

However, the unconditional distribution of *x*, which will necessarily be dependent on *y*, is not given a priori. Furthermore, the above condition gives no prescription on how to consistently and randomly construct appropriate objects, and does not reveal how the properties of these objects reflect on the character of dependence.

This can be remedied by deriving a criterion for dependence which is agnostic about the unconditional distribution of *x*, because from the perspective of data synthesis *P*(*x|y*) could be a matrix defined independently of *P*(*y*), which would then induce *P*(*x* ∩ *y*) along with *P*(*x*) through the chain rule relationship. A well-defined rule, consistent with this criterion, should (i) aid in the automation of generation of conditional distributions, (ii) reduce the number of parameters needed to synthesize a random network to the number of nodes and their arity, and (iii) allow to traverse arbitrarily defined collections of random networks with relative ease (by direct sampling).

Consider the following observation

### Lemma 1.

*Let x be a random variable with n outcomes, then the columns of conditional distribution P*(*x|y*) *are members of* (*n* – 1)-*simplex* 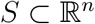.

*Proof*. Trivially, the elements of any column of a conditional distribution *p*(*x|y*) sum up to 1. Hence, because the variable *x* has *n* outcomes, all columns are members of the set defined by

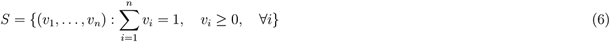

which is an *n*-simplex, and a subset of 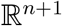

To simplify the notation and to emphasize that the conditional distribution *P*(*x|y*) can be defined independently, let *D* be an *n* × *m* linear operator whose columns are members of the (*n* – 1)-simplex *S*, so that *P*(*x*) = *D* · *P*(*y*) given some *P*(*y*).

### Lemma 2.

*If the column rank of D is at least two, then x defined by P*(*x*) = *D* · *P*(*y*) *is conditionally dependent on y*.

*Proof*. Suppose the rank of *D* is at least two, then there is at least one linearly independent column, i.e.

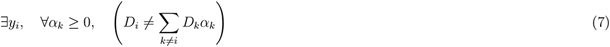

where *D_i_* is the *i*-th column of *D*. This implies that if for every index *k* ≠ *i* the coefficients are given by 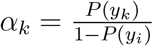, then

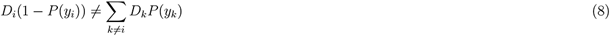

Rearranging the terms yields

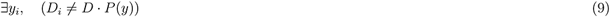

which is a restatement of the dependence criterion. Hence, there is at least one dependent event, making *x* dependent on *y*.

This lemma already guarantees that any matrix of appropriate size with rank of at least two should be sufficient to generate a conditionally dependent pair of variables. However, this does not elucidate additional constraints that an arbitrary conditional dependence may impose, and it does little to provide control over the structure of a synthetically constructed dependence relationship.

The following lemma addresses these shortcomings:

### Lemma 3.

*Let x be a random variable with n outcomes, dependent on a random variable y with m outcomes; then*

- *the convex hull C formed by the columns of D has at least two vertices;*
- *rank of D must be at least two;*
- *P*(*x*) *lies in the convex hull C formed by the columns of D;*
- *the number of vertices of convex hull C corresponds to the number of conditionally dependent events*.

*Proof*. Suppose *x* is conditionally dependent on *y*, then

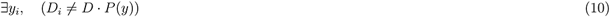

where *D_i_* is the *i*-th column of *D*, because *P*(*x*) = *D* · *P*(*y*). Rearranging the terms gives

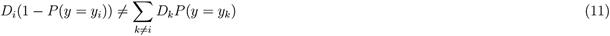

for some arbitrary *P*(*y*). Assuming that *P*(*y* = *y_i_*) ≠ 1, this yields

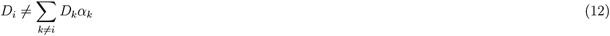

with coefficients defined on the probability simplex characterized by

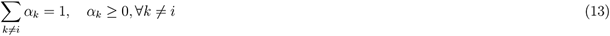

In other words, for *x* to be conditionally dependent on *y* for any *D* defined independently of *P*(*y*), there must be at least one column in *D* that must not be a convex combination of other columns. Therefore, this column is a vertex of a convex hull *C* ⊂ *S* formed by all convex combinations of columns in *D* characterized by

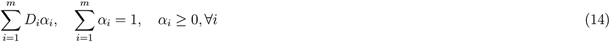

The above implies the following:

- the convex hull contains more than one member, and, therefore, has at least two vertices;
- because there are at least two vertices, this polytope must be embedded in at least a (2-1)-simplex, and, therefore, in at least 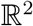 - hence the linear operator *D* must be at least of rank two;
- the unconditional distribution *P*(*x*) always lies in this convex hull, because *P*(*x*) = *D* · *P*(*y*) for any independently defined *D* and *P*(*y*);
- every conditionally dependent event of *y* corresponds to a vertex of the convex hull that envelops the columns of *D*.

Some of the immediately useful consequences of the above can be summarized in the following corollary:

### Corollary 1.

*Let x be a random variable with n outcomes and y be a random variable with m outcomes and D a linear operator characterized by P*(*x*) = *D* · *P*(*y*), *then*

- *x is conditionally dependent on y if and only if the rank of the linear operator D is at least two*.
- *Let d be the number of dependent ancestor events that the linear operator D of the size n* × *m and rank* = *r encodes; then r* ≤ *d* ≤ *m*.

Conditional distributions of higher ancestral complexity, i.e. *P*(*x*|∩*Y_k_*), can be treated as a pairwise distribution *P*(*x|y*) if we observe that ∩ *Y_k_* can be viewed as a single variable over the combinatorically prescribed joint events. In this situation, the only specification that is different from the already considered scenario with the pairwise conditional distribution is that the joint events are not given a priori. Rather, the joint events have to be derived as a function of the number and arity of ancestor variables in the DAG. For example, for two binary ancestor variables, there will be four possible events.

These results suggest an explicit approach for constructing random conditional dependence distributions (and generating synthetic data data from them) in a formally sound way, while ensuring that all relevant relationship properties are accounted for:

First, it should be noted that the numerical structure of the linear operator *D* is secondary to the properties of the convex hull formed by its columns, because it reflects the qualitative structure of the event space of the conditional dependence relationship. In particular, manipulating the properties of this convex polytope provides a way to choose the proportion of dependent and independent events in the event space.

Second, we observe that the columns of *D* must be sampled from the (*n* – 1)-simplex *S*. If sampled uniformly, this is equivalent to sampling from a Dirichlet distribution **Dir**(*α*) with the appropriate concentration parameters. However, our primary objective — constructing random dependence relationships — is equivalent to sampling polytopes of different shapes, sizes and locations from the simplex S, something that uniform sampling of individual simplex elements is not able to address. This can be detrimental to our purposes, as will become clear presently:

### Proposition 1.

*Let **t,p,q** be i.i.d. random variables obtained from **Dir*** (***α***) *with **α*** = (1,1), *and let D be a* 2 × 2 *matrix with columns **p** and **q**. Then the distribution of **z*** = *D* · *t is non-uniform*.

*Proof*. Note that all the variables including **z** are members of (2 – 1)-simplex, so all variables have two components, i.e. ***z*** = (*z*, 1 – *z*), ***t*** = (*t*, 1 – *t*), ***p*** = (*p*, 1 – *p*) and ***q*** = (*q*, 1 – *q*). Then ***z*** = *D* · ***t*** reduces to the single independent equation relating the components, namely *z* = *tp* + (1 – *t*)*q*. Using the substitution 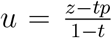 to reparametrize the integral yields the following expression for the cumulative distribution

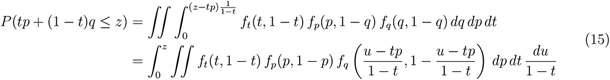

The corresponding density is obtained by differentiating with respect to *z* and evaluating the resulting integrals with the appropriate integration bounds. Using the requirement that 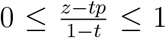 allows to infer the integration set for *p* as

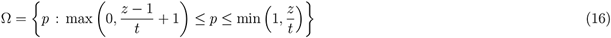

Applying the above to the evaluation of the integral yields

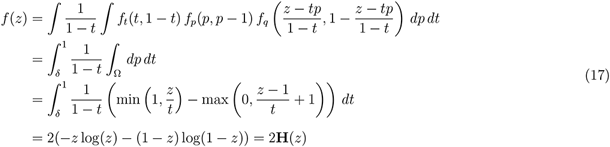

where **H** is the information-theoretic entropy. Hence, the resulting density is non-uniform.

The above implies that the naively constructed *D* induces a non-uniform distribution, strongly favouring simplex elements with higher entropy. Furthermore, as the number of columns in *D* grows, the components of ***z*** = *D* · ***t***, viewed as a weighted sum of the columns of *D*, satisfy the conditions of Lyapunov Central Limit Theorem. To see this consider the following simple case:

### Proposition 2.

*Let D_i_, a member of* 2 – 1-*simplex, the i-th column of a* 2 × *n matrix D, and **t**, a member of m* – 1-*simplex, be random variables obtained from **Dir*** (***α***) *with **α*** = (1,1) *and an n-tuple **α*** = (1,…, 1) *respectively. Then each of the components of **z*** = *D*·***t*** *tends towards a normal distribution*.

*Proof*. First note that the expectation, variance and density of the *i*-th component *t_i_* of ***t*** obtained from ***Dir***(***α***), with *α_k_* = 1 and 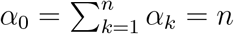, are given by

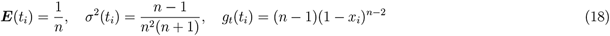

For *d_i_*, the density is *g_d_*(*d_i_*) = 1. Next, let *z_i_* = *t_i_d_i_* and *y_i_* = *z_i_* – ***E***(*z_i_*), where 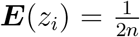. Using the binomial expansion, the density for *z_i_* can be obtained as follows:

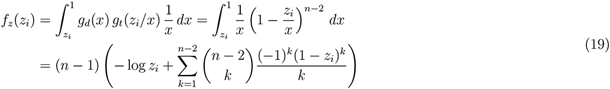

Further, define

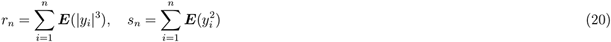

Since the expression for any order moment of *y_i_* is the same for any *i*, rewrite the above as

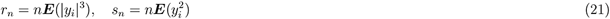

Applying the Lyapunov inequality to the third absolute moment yields the bound

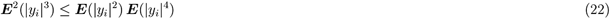

Noting that

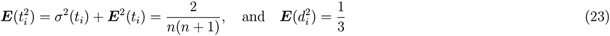

allows to evaluate

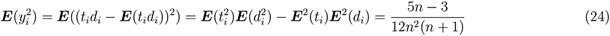

Using the binomial expansion to evaluate the forth moment yields

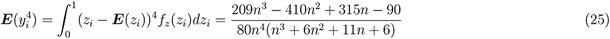

Then the Lyapunov condition combined with the bound obtained above yields

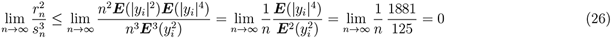

Therefore, as *n* → ∞

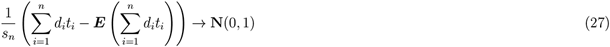

in distribution.

Clearly, with naively constructed conditional probability *D*, the distributions downstream of the root node in the network will tend towards the maximum entropy states — this tendency further amplified down the ancestor/offspring chains. This is undesirable, as it will significantly shrink the range of accessible models. Additionally, there is no reason to believe that the situation will be any different for *D* of arbitrary size, or that the slow convergence rate in Proposition 2 would alleviate this effect, since *n* grows exponentially with the number of ancestors.

The above analysis reinforces the idea that, in order to uniformly cover the model space, we would require explicit specification for a class of distributions over simplicies that will allow us to construct polytopes of different shapes, sizes, and locations. While the detailed analysis/prescription is outside the scope of this study, and will be pursued elsewhere, here we suggest that the desired behavior can be achieved by manipulating concentration parameters ***α*** in ***Dir***(***α***). However, the observed correspondence between the geometrical object encapsulated in the linear operator *P*(*x|y*) and the conditional dependence relationship that it represents, already fully addresses the concerns raised above. Any further developments — such as uniform sampling of polytopes from the encompassing simplex — are not strictly necessary at this stage, because our primary goal is not to generate the “proper” typical linear operators associated with conditional dependency, but rather to provide complete justification of, and technical basis for, the data synthesis method, wherein we start with the unconditional distribution of the set of root nodes and sequentially generate the full joint probability distribution of the network given any arbitrary DAG structure. Note that the DAG structure is equivalent to an adjacency (0,1)-matrix where the acyclicity requirements necessarily dictates triangular shape and zero main diagonal. Hence, to produce arbitrary DAG structures, it is sufficient to generate random triangular adjacency matrices.

Interestingly, the seemingly relaxed and unimposing nature of the minimum requirement for dependence (stated in the Corollary 1) further reinforces the notion (discussed in the preceding section) that probabilistic relationships are, by their very nature, “generic” / implicit.

In conclusion, after defining all the necessary parameters, the sample construction can be carried out starting with the sampling of the set of root nodes, and proceeding sequentially forward following the rule that, given a particular ancestral sample *y*\*, the joint probability distribution for the downstream node combined with the ancestral sample is given by

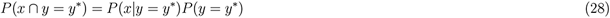

i.e. for each sample *y* = *y*\* the sample *x* = *x*\* is obtained with probability *P*(*x* = *x*\*|*y* = *y*\*). This serialized construction of samples will produce a single row of data with values corresponding to nodes in the probabilistic DAG. This row will then constitute a single sample from the given DAG. Repeating this procedure as necessary will generate a dataset of prescribed size. We discuss sample size considerations below in Results: Section 5.

## 3 Methods, Proofs and Results: Random graph distribution

To estimate the error of the structure recovery for the inverse solver (i.e., the actual BN reconstruction algorithm), the forward solver (the synthetic data generator) needs to not only synthesize data from a given model, but also generate the random model structures (joint probability factorizations) that the data is sampled from. This can be realized by producing random DAG adjacency matrices, or, more specifically, triangular extractions from symmetric adjacency matrices to account for the acyclicity of structures.

Thus, synthesized random structures should be distributed in a prescribed manner, a crucial characteristic without which error estimation will necessarily lead to unreliable results. Therefore, we need to establish some of the basic properties of DAG structure distributions.

The discrete nature of graph configurations implies nonuniform distribution across various graph densities for combinatorical reasons. For a graph of *n* variables with a given topological ordering the maximum density (maximum number of edges) possible is given by

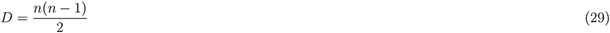

Then the total count *C* of possible graph configurations with a density *d* < *D* is given by the binomial coefficient

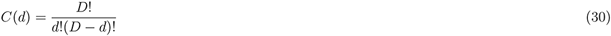

which implies that structures with density around 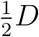 are more numerous than symmetrically higher or lower density structures. The total number of possible configurations of any density is

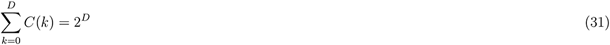

Let *δ* be the density valued function over graph structures. Then the probability that a graph 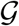 has density 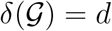 is

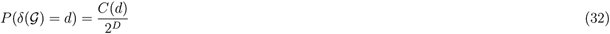

The joint probability of obtaining a configuration 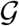 with density *d* is therefore

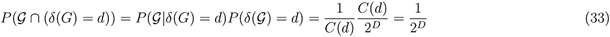

where the conditional probability of 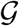 given the prescribed density is

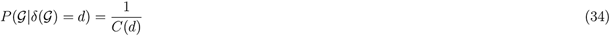

Of particular interest is the expression for 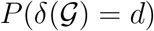 because it implies that structures of medium density are more likely than any other. This effect introduces bias in error estimation, and would compromise the performance of a naive stochastic structure recovery method that searches the state space by evaluation only, not accounting for the fact that medium density states will occur more frequently.

Technically, this could be remedied by a forced uniform sampling across various density groups, thereby increasing the sampling rate over lower and higher density groups relative to the medium density groups or, similarly, by decreasing the sampling rate over the medium density groups. However, this may be impractical, because even for a problem of moderate size, the number of possible configurations at the extremes of the distribution is sharply limited relative to the medium density graphs, which would make it virtually impossible to maintain a constant sampling error. For a 16-node structure the above translates to 120 configurations with one edge, 840261910995 configurations with 8 edges, and on the order of 10^34^ medium-density configurations with 60 edges. Therefore, if a 10% sampling rate is to be maintained at any cost, then the number of samples across all density groups is in the order of 10^35^ (with most of the computational load in the medium-density groups). Therefore, we conclude that in practice, instead of pursuing a uniform sampling error, one should aim to address this situation analytically.

## 4 Methods, Proofs and Results: Forward inference over Markov Blankets

It is important to assure the congruence of independence / probabilistic inference mechanisms in the simulation framework (as described above) and during the BN construction process. In this section, we will dissect the ‘localized” probabilistic inference within a BN by utilizing the concept of Markov Blanket. By definition, a Markov Blanket *M* of a particular variable *x* renders such ‘center” variable conditionally independent from the rest of the BN (i.e. BN sans the Markov Blanket), with the ‘‘periphery” variables of *M* completely determining the state of the center variable assuming their states are known. This naturally leads to the problem of estimating conditional probability of *x* as a function of the states of periphery variables in *M* – *x*.

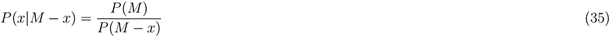

The computational difficulties here are the same as with the estimation of any sufficiently complex joint probability. Often we would lose accuracy due to the sampling error before we can estimate most joint events unless all the variables are defined analytically, and the dimensionality of the computation for such an estimation grows superexponentially with the number of variables. However, these difficulties can be mitigated by relying on the structural information encoded in the BN. If correct, the simplifications introduced into factorization sufficiently reduce the computational burden and the sampling error, thereby making the desired estimation not only feasible, but also more reliable. And although some of the difficulties alleviated are traded for the multiplicative propagation of error, the end result is still preferable to the direct loss of resolution.

Let *M* be Markov Blanket of *x* and let *a*(*z*) ⊂ *M* denote the ancestor set of *z* ∈ *M* in *M* characterized by the following statement

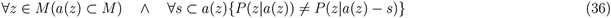

where *s* is any subset of the ancestor set. Further, let *d*(*z*) ⊂ *M* be the descendant set of *z* ∈ *M* characterized by the following

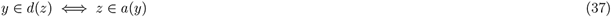

And, for notational convenience, let

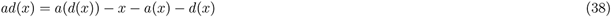

so that *ad*(*x*) represents the set of ancestors of members of *d*(*x*) excluding *x* itself that are neither direct ancestors or descendants of *x* themselves, i.e.

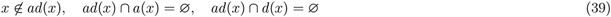

Then the joint probability of MB of *x* can be factored as follows

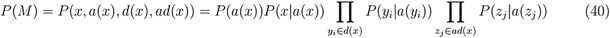

The above equation can be derived through the following scheme

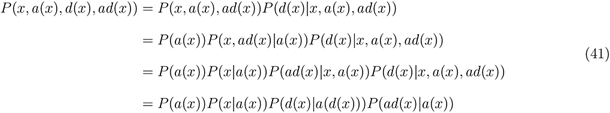

Methodical application of the chain rule to the remaining complex conditional probability terms yields the result in (40). Also, note that

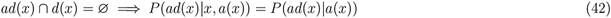

and that

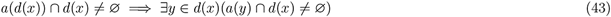

Equation (40) provides direct access to all joint probabilities for any instantiation of variables in *M*, allowing us to proceed with the estimation of the desired conditional probability

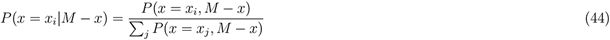

marginalizing over *x* in the denominator for any given state *x_i_* of *x* in the numerator. Undoubtedly, the division operation itself introduces further numerical error into the estimate, but the classical numerical techniques are applicable in this situation.

Different types of inference are possible under the same assumptions, utilizing the same set of equations. For example, one could assess the probability of a periphery variable *z* ∈ *M* being in a state *z_i_* in order for the center variable *x* ∈ *M* to be observed in a state *x_j_*:

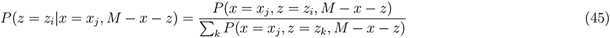

In essence, the above type of inquiry allows to make a judgment about the most likely scenario that coincides with the *x* = *x_j_* event, and this inquiry could be extended to more than one periphery variable.

Furthermore, the idea that periphery variables can predict the state of the center variable with a prescribed accuracy can also be utilized in structural optimization during the BN structure search. Local prediction accuracy as an objective optimization criterion is inherently practical because it attempts to maximize the utility of the result rather than concentrating exclusively on a set of more abstract notions associated with information-theoretic properties.

## 5 Results: Sample size considerations

In practice, it is important to evaluate how many samples should be generated (as prescribed in Section 2) for the estimation of the joint probability to be sufficiently robust. As a representative example, let us consider a network of eight variables, with arity of three and average connection density of 0.8. This network can assume 3^8^ = 6561 possible configurations, and it is straightforward to analytically compute the probabilities of all configurations. Subsequently, we can evaluate the effect of sample size in estimating the “true” distribution. Figure 1 shows the analytical and estimated distribution for all possible configurations in our example, for four (increasing) sample size values. To quantify the differences between the distributions, we used Jensen-Shannon divergence (JSD), a distributional divergence metric (Figure 2). These results confirm that our procedures converge to zero distributional divergence with the increasing sample size.

**Figure 1:**
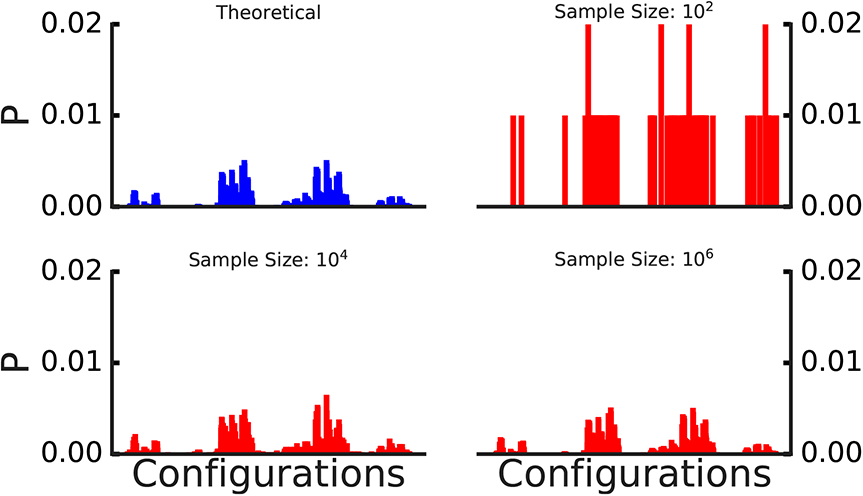
Comparison of the analytical and estimated sampled distributions of all possible configurations of a given 8-nodes random graph with average variable connection density and arity being 0.8 and 3.0, respectively. The analytical distribution is shown in top left panel (blue); distributions estimated using sample size (10^2^, 10^4^, 10^6^) are shown in red.

**Figure 2:**
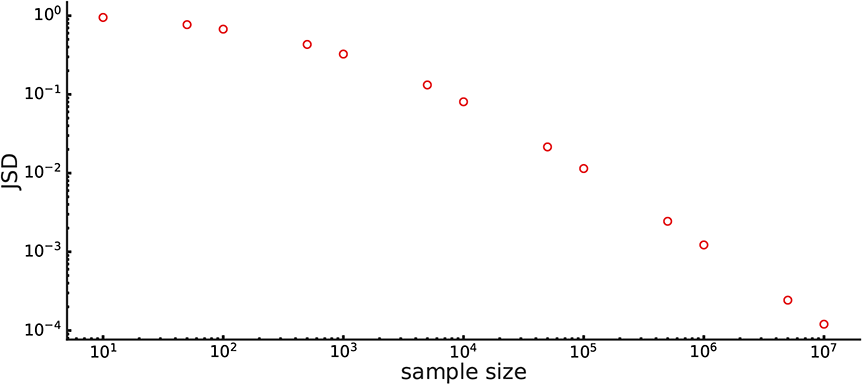
Jensen-Shannon divergence (JSD) between the analytical and estimated sampled distributions of all possible configurations of a given 8-nodes random graph with average variable connection density and arity being 0.8 and 3.0, respectively. JSD, shown as a function of sample size, is obtained by averaging 100 replicas for each sample size value.

Given that the analytical joint distribution over the network is calculated during the simulation, the sample size necessary to achieve the desired representational accuracy can be estimated via the multinomial distribution. If ***p*** = (*p*_1_,…, *p_k_*) is the density of the joint events, and ***x*** = (*x*_1_,…, *x_k_*) are the sample counts of the observed joint events, then the probability of sampling a particular sequence of counts ***x*** is given by the multinomial *f*(***x, p***). This allows to formulate a number of possible accuracy constraints. For example, controlling the maximum deviation from the idealized count *N_**p**_*, with *N* = Σ_*i*_ *x_i_*, can be achieved by estimating N necessary to bring the probability of such a maximum deviation bellow a certain threshold e, i.e. *P*(max_*i*_|*N_p_i__* – *x_i_*| > *δ*) < *ϵ*, so that for a defined accuracy level *δ* all deviations beyond that have a small probability of occurrence.

## 6 Discussion and Conclusions

Simulation studies based on synthetic data generation are ubiquitous in the machine learning / computer science domain. They are instrumental in objectively assessing the performance of descriptive/predictive modeling methods, such as BNs. However, comprehensive and realistic simulations frameworks are comparatively underdeveloped in the context of network-centered systems biology methodology. Most of the BN simulation studies rely on SEM approximations — not necessarily a good fit with biological reality. In this study, we presented theoretical and methodological considerations that can be used to develop a more realistic, fully probabilistic, synthetic data generation framework for biological BNs.

At this time, we have developed algorithms and software for generating synthetic data for randomly generated discrete-variables probabilistic DAGs. These are included as part of our “BNOmics” BN modeling software package (Gogoshin et al. (2017)), which is freely available from the authors, as well as from the bitbucket open source distributary (https://bitbucket.org/77D/bnomics). In the future, we intend to expand our algorithms to take into account mixed data types (continuous/discrete variable mix, specifically).

Two outstanding issues, relevant to both the BN reconstruction, and synthetic data generation / BN performance evaluation, are (i) whether the Markov Blankets (within the BNs) can in principle be recovered with consistency (provided that the “forward generation” and “backward reconstruction” processes are sufficiently aligned, as in our proposed framework), and (ii) whether it is possible to incorporate a strict, rigorous definition of dependence / conditional independence in the BN reconstruction process, as opposed to concentrating on establishing causation from correlation. We are cautiously optimistic on both counts, but more investigation is needed, and will be forthcoming.

Much of the work detailed in this communication was spurred by our ongoing collaborative immunooncology study that involves comparative BN analyses of multi-dimensional FACS (fluorescence-activated cell sorting) and other immuno-oncology datasets obtained from patients with gastrointestinal and breast cancer undergoing immunotherapy treatments. In that study, we aim to construct and compare/contrast BNs representing immune signaling network states before and after therapy, in responders and nonresponders to therapy. In the future, we plan to generate synthetic datasets that more closely align with the real, observed FACS and other immuno-oncology datasets — this will allow us to rigorously validate the reconstructed immune signaling BNs and corresponding biological results. Based on our experience working with such data, it is our opinion that each and every BN analysis should, ideally, be accompanied by a corresponding simulation study built around simulated data that is as close as realistically possible to the actual biological data under consideration.

## 7 Acknowledgements

The authors are grateful to Arthur D. Riggs, Russell C. Rockne, Peter P. Lee and Amanda J. Myers for many stimulating discussions and useful suggestions on the applicability of BN methodology to a wide variety of biomedical data. Research reported in this publication was supported by the National Institutes of Health under grant numbers P30CA033572, U01CA232216 and R01LM013138. Additional support by the Susumu Ohno Chair in Theoretical and Computational Biology (held by A.S.R.) and the Susumu Ohno Distinguished Investigator fellowship (to G.G.) is kindly acknowledged. The funders had no role in study design, data collection and analysis, decision to publish, or preparation of the manuscript. The content is solely the responsibility of the authors and does not necessarily represent the official views of the National Institutes of Health.

## 9 Data Availability Statement

Relevant code and software are available directly from the authors, or as a part of the BNOmics package, at https://bitbucket.org/77D/bnomics.

